# Protein Co-Evolution Strategies Detect Predicted Functional Interaction Between the Serotonin 5-HT_2A_ and 5-HT_2C_ Receptors

**DOI:** 10.1101/512558

**Authors:** Bernard Fongang, Kathryn A. Cunningham, Maga Rowicka, Andrzej Kudlicki

## Abstract

Serotonin is a neurotransmitter that plays a role in regulating activities such as sleep, appetite, mood and substance abuse disorders; serotonin receptors 5-HT_2A_R and 5-HT_2C_R are active within pathways associated with substance abuse. It has been suggested that 5-HT_2A_R and 5-HT_2C_R may form a dimer that affects behavioral processes. Here we study the coevolution of residues in 5-HT_2A_R and 5-HT_2C_R to identify potential interactions between residues in both proteins. Coevolution studies can detect protein interactions, and since the thus uncovered interactions are subject to evolutionary pressure, they are likely functional. We assessed the significance of the 5-HT_2A_R/5-HT_2C_R interactions using randomized phylogenetic trees and found the interaction significant (p-value = 0.01). We also discuss how co-expression of the receptors suggests the predicted interaction is functional. Finally, we analyze how several single nucleotide polymorphisms for the 5-HT_2A_R and 5-HT_2C_R genes affect their interaction. Our findings are the first to characterize the binding interface of 5-HT2AR/5-HT2CR and indicate a correlation between this interface and location of SNPs in both proteins.

## INTRODUCTION

Serotonin (5-hydroxytryptamine, 5-HT) is a monoamine neurotransmitter with broad regulatory roles within the central and peripheral nervous systems with complex actions transduced by 5-HT binding to one ligand-gated cation channel and 13 G protein-coupled receptors (GPCRs). Of the 5-HT GPCRs, the 5-HT_2_ receptor family includes the 5-HT_2A_ receptor (5-HT_2A_R), 5-HT_2B_R and 5-HT_2C_R on the basis of their structural and operational characteristics (Hannon and Hoyer 2008). The brain is particularly enriched in expression of the 5-HT_2A_R and 5-HT_2C_R that govern normal behavioral (e.g., appetite, mood, sleep) and physiological function (e.g., circadian rhythm, endocrine secretion). Furthermore, dysfunctional 5-HT_2A_R and 5-HT_2C_R systems are implicated in a broad range of neuropsychiatric disorders (e.g., addiction, depression, schizophrenia) and obesity (Hamon and Blier 2013)(Howell and Cunningham 2015). These receptors couple to Gα_q/11_ to activate the enzyme phospholipase Cβ which generates intracellular second messengers inositol-1,4,5-trisphosphate and diacylglycerol, leading to increased calcium release from intracellular stores (Hannon and Hoyer 2008). The homomeric form of the 5-HT_2A_R (Iglesias, Lage et al. 2016, Iglesias, Cimadevila et al. 2017, Iglesias, Cimadevila et al. 2017) and 5-HT_2C_R (Herrick-Davis, Grinde et al. 2012, Herrick-Davis, Grinde et al. 2015) is proposed to be main signaling unit, while both are reported to form functional heterodimeric complexes with other GPCRs (Albizu, Balestre et al. 2006, Albizu, Moreno et al. 2010) (Schellekens, Dinan et al. 2013, Schellekens, van Oeffelen et al. 2013). Employing co-immunoprecipitation (Co-IP) assays, we recently demonstrated that the two receptors form a protein complex in rodent brain, expression of which tracked with phenotypic behaviors (Anastasio, Stutz et al. 2015) while bioluminescence resonance energy transfer (BRET) assays detected a very close (within 10 nM) interaction in transfected cells (Felsing et al., in press; (Moutkine, Quentin et al. 2017). The 5-HT_2A_R:5-HT_2C_R protein complex triggers intracellular signaling in live cultured cells that is uniquely distinct from each protomer alone (Moutkine, Quentin et al. 2017). Further studies are required to predict biological features of relevance such as residue contacts within the 5-HT_2A_R:5-HT_2C_R protein structures and protein functional sites.

Several experimental techniques are useful in the study of protein dimerization, including Co-IP, BRET, tandem affinity purification with mass spectrometry, yeast two-hybrid screens, protein-fragment complementation assays, NMR spectroscopy and split luciferase complementation assays (Rigaut, Shevchenko et al. 1999, Uetz, Giot et al. 2000, Sekar and Periasamy 2003, Keene, Komisarow et al. 2006, Rochette, Diss et al. 2015). Although useful in the detection of dimerization interfaces, these methods cannot distinguish between direct and indirect interactions, are frequently performed under non-physiological conditions, and remain time consuming and costly. Moreover, each approach probes a different aspect of the interaction, and suffers from the potential for assay-specific experimental errors; also, experimental approaches may miss transient interactions crucial to biological processes, limiting their utility in providing definitive answers as to whether an interaction is functional *in vivo* (Phizicky and Fields 1995). An alternative approach for identifying functionally significant protein-protein interactions is through the study of evolutionary coupling, or coevolution of residues in these proteins along the phylogenetic tree (Halabi, Rivoire et al. 2009, Marks, Hopf et al. 2012, de Juan, Pazos et al. 2013, Hopf, Scharfe et al. 2014, dos Santos, Morcos et al. 2015, Gueudre, Baldassi et al. 2016). The concept underlying evolutionary coupling is that, to preserve function, a mutation in one interacting residue is likely to be compensated by a complementary mutation in the other (for review) (de Juan, Pazos et al. 2013). The key advantage of this approach is that interactions between residues are detected not only based on their physical proximity, but upon evolutionary pressure, and therefore are more likely to be functionally relevant for biology. Such methods have detected transient interactions, including allosteric changes in proteins (Lockless and Ranganathan 1999)(Hatley, Lockless et al. 2003, Suel, Lockless et al. 2003)), as long as they are important for the protein function. For almost two decades, co-evolution of residues in protein sequences has been valuable for inferring residue-residue interactions (contacts) within small, bacterial proteins with large numbers of homologs (Lockless and Ranganathan 1999, Morcos, Pagnani et al. 2011, Ochoa, Garcia-Gutierrez et al. 2013). Several statistical approaches to studying co-evolution have been developed (Morcos, Pagnani et al. 2011, Marks, Hopf et al. 2012, Sulkowska, Morcos et al. 2012, Ekeberg, Hartonen et al. 2014, Hopf, Scharfe et al. 2014), including direct coupling analysis (DCA) (Morcos, Pagnani et al. 2011). With the rapid increase in the number of sequenced animal genomes, these methods have become feasible for assessments of protein interactions in metazoans, including humans, and for larger proteins and even protein complexes (Hopf, Colwell et al. 2012, Naveed, Xu et al. 2012, Hopf, Scharfe et al. 2014, Kim, DiMaio et al. 2014, Ovchinnikov, Kamisetty et al. 2014, dos Santos, Morcos et al. 2015, Tesileanu, Colwell et al. 2015, Abriata, Bovigny et al. 2016, Champeimont, Laine et al. 2016, Feinauer, Szurmant et al. 2016, Lua, Wilson et al. 2016, Neuwald 2016, Yu, Vavrusa et al. 2016). Here, we adapted DCA to studying interactions between the 5-HT_2A_R and 5-HT_2C_R, which are particularly challenging due to their size, α-helical transmembrane domain structure, and limited structural resolution for these proteins. Next, we utilized simulated phylogenetic trees to demonstrate that the identified interactions are indeed statistically significant. We interpret the results in the context of the structure of the 5-HT_2A_R and 5-HT_2C_R proteins, as well as the known polymorphisms in the genes coding for these proteins. Finally, we present additional evidence supporting the 5-HT_2A_R and 5-HT_2C_R interaction hypothesis, based on temporal co-expression of the receptor genes in the primate circadian cycle. We will also discuss how our method can be adapted and extended to other challenging classes of protein interactions, such as interactions between other G-protein coupled receptors that are targets of 30% of FDA approved medications (Filmore 2004, Overington, Al-Lazikani et al. 2006).

## RESULTS AND DISCUSSION

### Coevolution of residues in the putative 5-HT_2A_R/5-HT_2C_R heterodimer

Coevolution between residues of heterodimeric proteins can be computed using a wide range of methods from Mutual Information (MI), Statistical Coupling Analysis (SCA), to the Direct Coupling Analysis (DCA).

Here, we chose the DCA algorithm (Morcos, Pagnani et al. 2011) among Evolutionary Coupling (EC) methods because of its superior ability to model the whole data set rather than individual pairs of residues, helping to distinguish between direct functional residue interactions and correlations resulting from indirect interactions. Sequences of 5-HT_2A_R and 5-HT_2C_R from 231 vertebrates were obtained from *GENBANK* (Benson, Lipman et al. 1993) and aligned using *Clustal-Ω* (Sievers, Wilm et al. 2011), resulting in a concatenated Multiple Sequence Alignments (cMSA) of the 5-HT_2A_R and 5-HT_2C_R.

We compared the cMSA with structural domain data and secondary structure prediction for the receptor proteins computed using stand-alone versions of PSIPRED (McGuffin, Bryson et al. 2000) (Methods). Both in 5-HT_2A_R and 5-HT_2C_R (Fig.S1), the transmembrane domains are highly conserved compared to the extra- and intra-cellular domains.

### The DCA map of 5-HT_2A_R and 5-HT_2C_R interactions

Evolutionary pressure between two proteins can be visualized as a N residues × M residue map (a DCA map), where each point represents the strength of the coupling between a pair of residues. The DCA map of the predicted residue interactions in the 5-HT_2A_R – 5-HT_2C_R heterodimer was generated with the top 5% of the *471 × 458* interactions (*471* and *458* are the lengths of 5-HT_2A_R – 5-HT_2C_R, respectively) (Fig. 2). For DCA to be successful, certain level of variation in residues in a given position is needed, completely conserved residues will not give any coevolution signal. In our case, such variability is sufficient in intra- and extracellular domains of 5-HT_2A_R and 5-HT_2C_R (supplementary Figure SF1), but their transmembrane domains are too well conserved (60% of transmembrane residues have a 100% conservation). Due to this high level of sequence conservation of the Transmembrane domains (*TM*), they do not produce strong signal, unlike IC and EC. This result should not be interpreted to me that there are no evolutionary conserved interactions involving TM domains, but that, due to high conservation of TM domains, their functional interactions are more challenging, or inaccessible, to studying using DCA methods.

**Figure 1:**
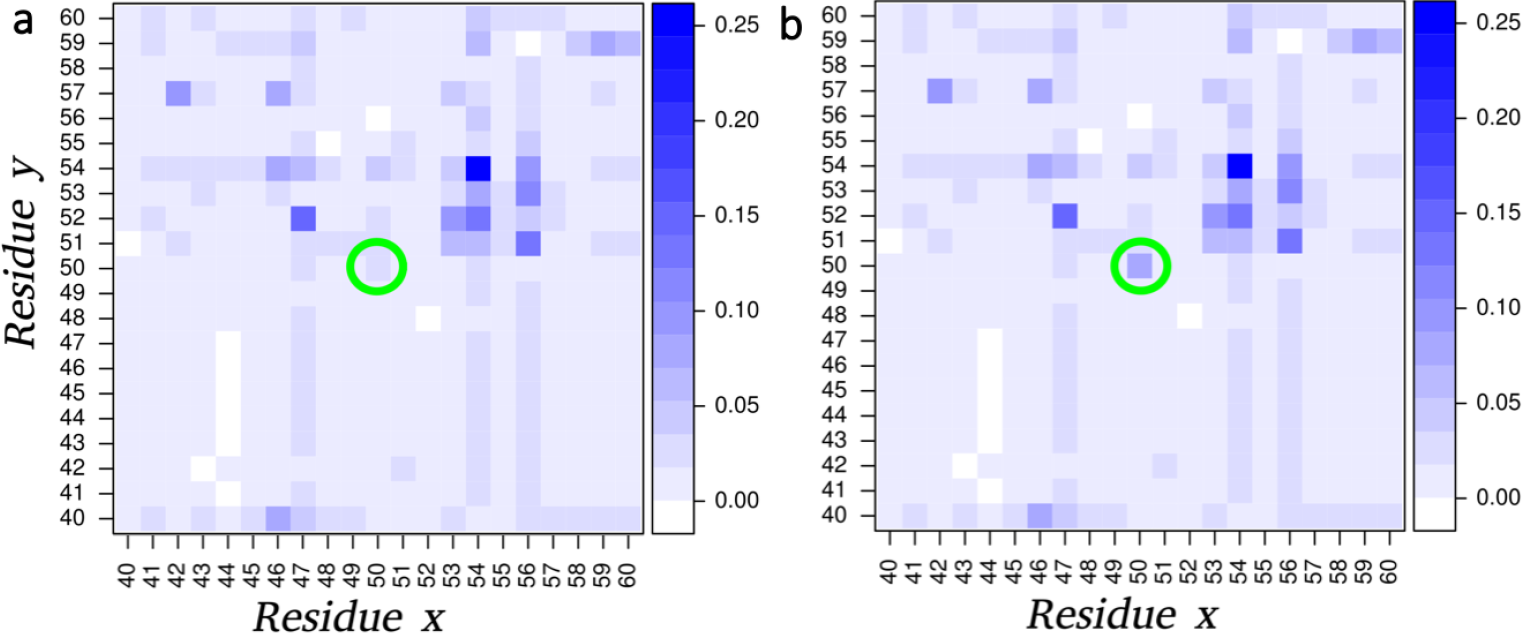
Convolved DCA signal. Neighborhood contribution to DCA Evolutionary score between residues. The original DCA signal (a) is convolved by a kernel centered around residues at position (50,50) on a window of 21 amino-acid. The final convolved signal (b) has been scaled to include the contribution of the neighboring residues.

**Figure 2:**
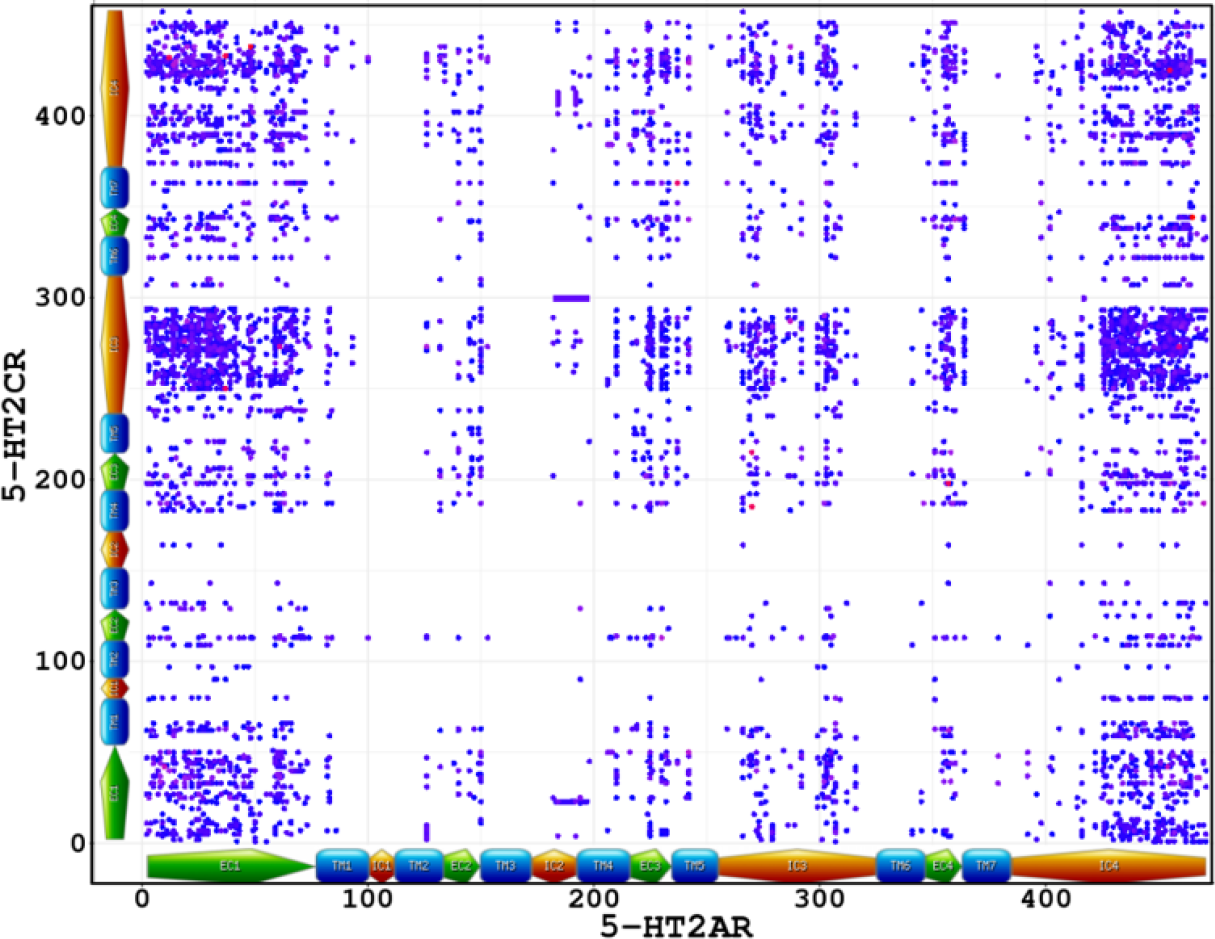
The raw, non-optimized DCA map of 5-HT_2A_R - 5-HT_2C_R. The figure shows top 5% of the interaction between the 2 receptors. Receptor domains are colored code (TM= Transmembrane, IC = Intra-Cellular, EC = Extra-Cellular). Due to high conservation of the TM domain, most signal are located at EC and IC domains (List of interacting sites in Supplementary Data File SDF2)

### Postprocessing the 5-HT_2A_R/5-HT_2C_R DCA map: Filtering out the false positives from the 5-HT_2A_R/5-HT_2C_R 2A-2C DCA map by Local convolution

Traditional DCA maps are interpreted in terms of strongest signal between residues, but since a strong signal in an island of background noise is not necessary an indication of dimerization, additional post-processing is needed to improve the inference of interacting peptides. Moreover, even though DCA is designed to assign higher scores to direct contacts than to indirect interactions, coevolution between indirectly interacting residues is still visible in the DCA maps (Figure 2), especially signal corresponding to allosteric interactions between distant residues. Such allosteric interactions may be responsible for the peaks connecting intracellular and extracellular domains. These interactions are likely mediated by transmembrane domains, and the lack of coupling signal in the transmembrane domains is due to the very high conservation of transmembrane residues that renders them inaccessible to detection with DCA.

Because DCA based methods are known to generate large amount of isolated False Positive (FP) contacts (Morcos, Pagnani et al. 2011, Sulkowska, Morcos et al. 2012, Ekeberg, Hartonen et al. 2014, dos Santos, Morcos et al. 2015), we have developed a method (see Methods section), based on local convolution of Evolutionary Coupling Scores (ECs) with a Gaussian kernel specific to the secondary structures of the protein, to reduce the FP and thus prioritize the most likely interacting peptides in protein complexes. The resulting filtered map of 5-HT_2A_R/5-HT_2C_R, for several values of the normalized variances and average length of interacting peptides, is shown in Figure 3. It appears that local convolution of ECs helps identify most likely interacting peptides (hot spots regions in Figure 3, panel d).

**Figure 3:**
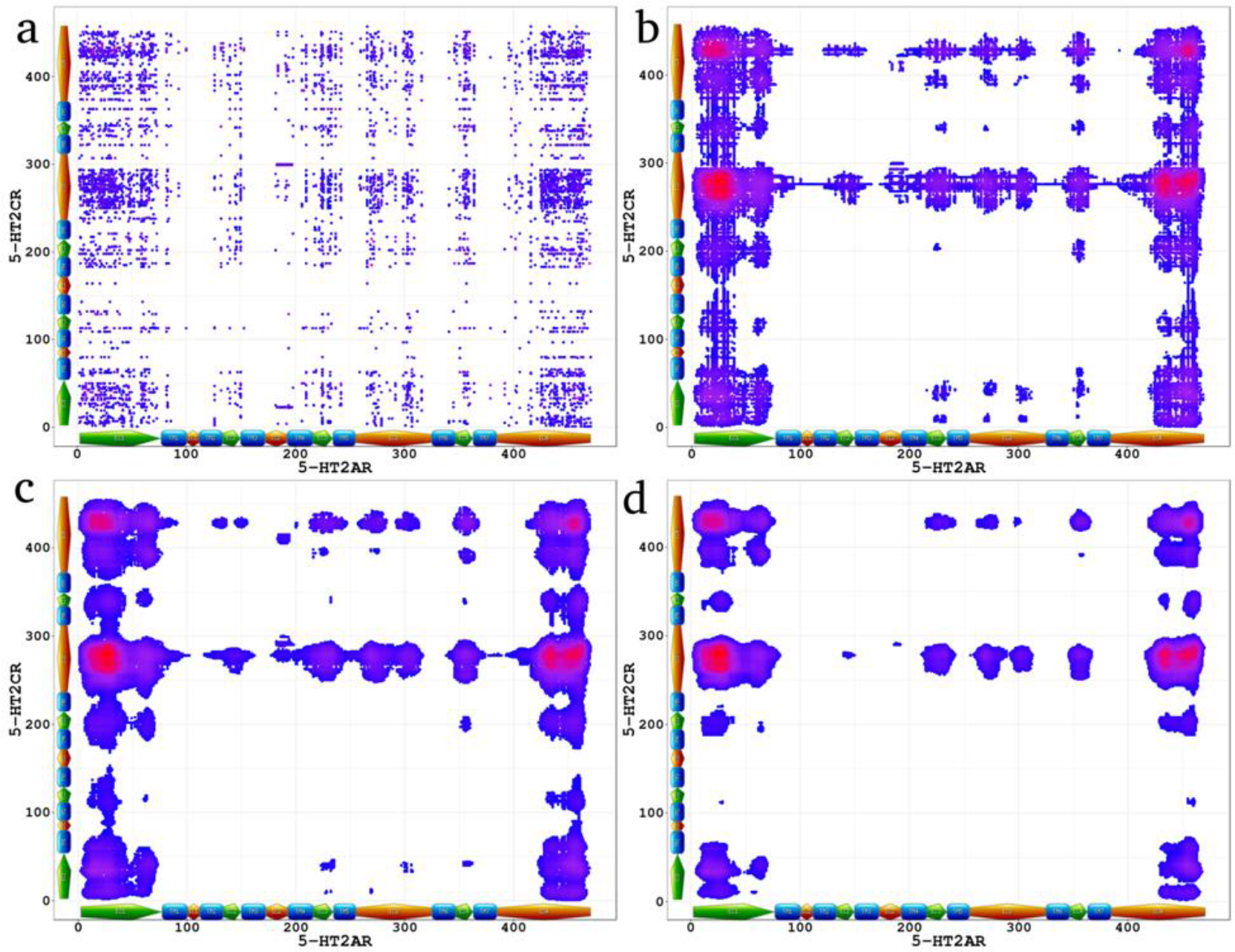
Convolved ECs of 5-HT_2A_R - 5-HT_2C_R highlights most likely interacting peptides. The local convolution method was applied to raw ECs (a) using optimized parameters as described in the Methods section (b to d).

### Protein domains involved in the predicted 5-HT_2A_R/5-HT_2C_R interaction

Because one of the main applications of the present study is to guide experiments determining the interacting peptides of oligomers, it is important to estimate the frequencies of domain-domain interactions which are the focus of experimental protocols such as point mutations. Both 5-HT_2A_R and 5-HT_2C_R consist of four Intra-Cellular domains (labelled IC1 to IC4), four Extra-Cellular domains (labelled EC1 to EC4), and seven Transmembrane domains (TM1 to TM7). For each individual domain in one receptor, we evaluated how frequently it interacts with other domains in adjacent protein. Only the top 5% interactions were considered. EC1 is the domain on 5-HT_2A_R with more significant interactions whereas IC3 on 5-HT_2C_R is the domain with more interactions. (Figure 4.a and Figure 5).

**Figure 4:**
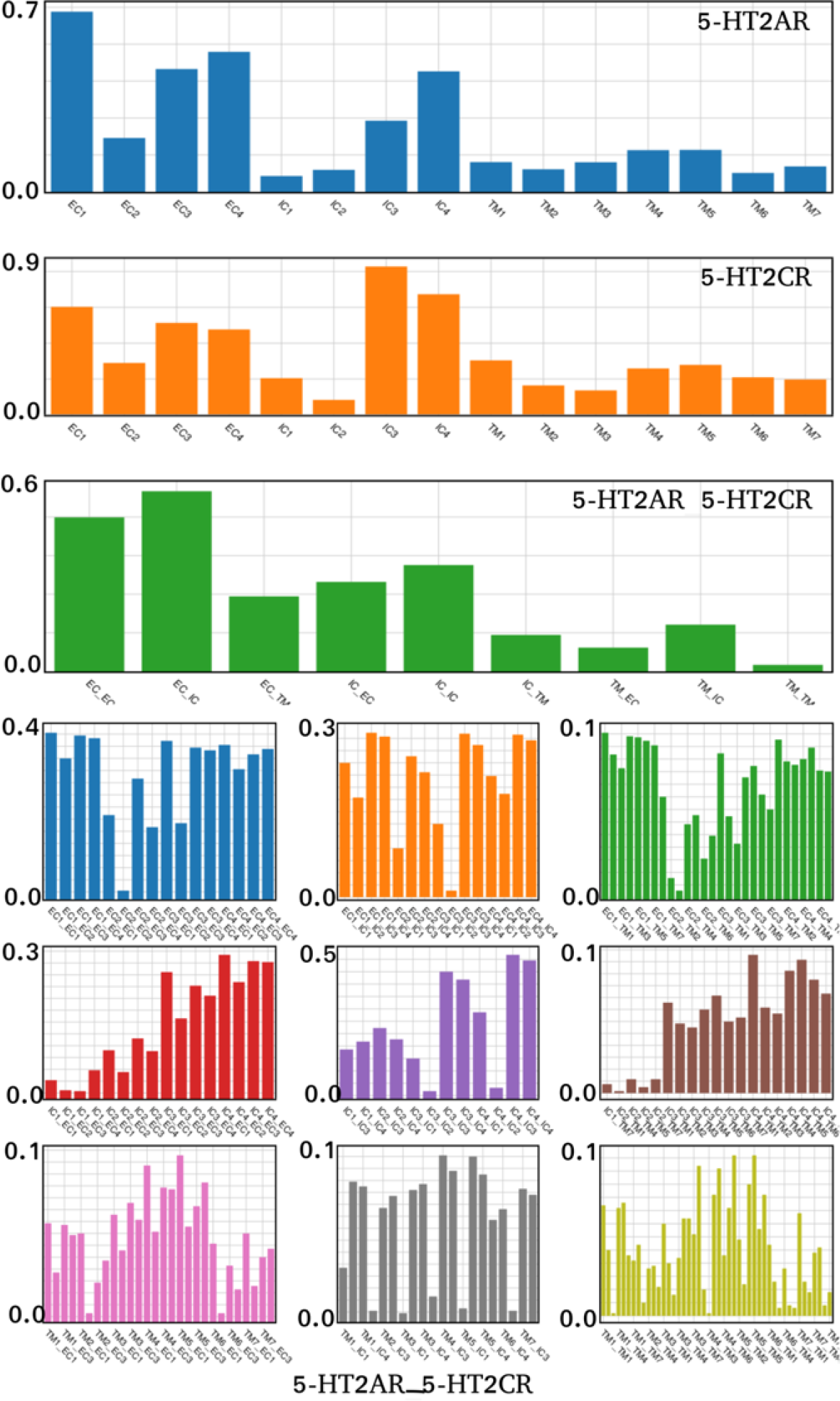
Domain interaction of 5-HT_2A_R - 5-HT_2C_R. We evaluate how frequently domains interact with each other (EC: Extra-cellular, IC: Intra-Cellular, TM: Transmembrane domain).

**Figure 5:**
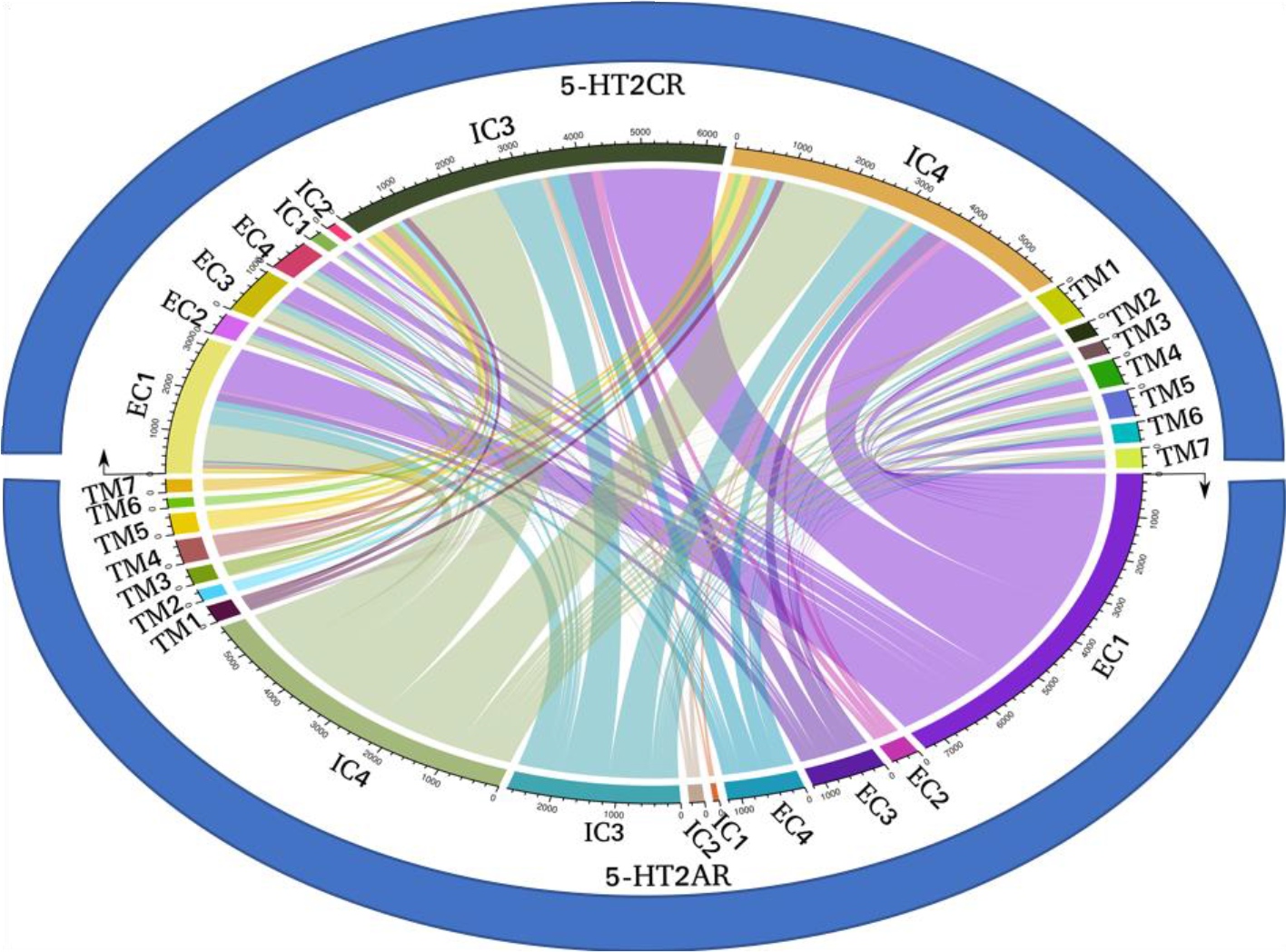
Domain interaction of 5-HT_2A_R - 5-HT_2C_R. A graph representing the interactions between the specific domains. Arc widths corresponds to numbers of inter-protein residue pairs with DCA signal greater than 0.2. (EC: Extra-cellular, IC: Intra-Cellular, TM: Transmembrane domain).

### Randomized phylogeny to assess the statistical significance of the heterodimerization

To compute the statistical significance of the heterodimerization, we used the randomized phylogeny which is defined as follows: in the joint MSA, we randomized the assignment of 5-HT_2C_R sequences to species while keeping the 5-HT_2A_R sequences unchanged and computed the ECs as previously described. For each randomized ECs, the number of predicted contacts higher than a threshold (minimum ECs score of the top 5% of the original ECs). Of the 100-randomized phylogeny ECs performed, only one predicted a number of contacts higher than the actual ECs suggesting the predicted interacting peptides between the two receptors is not random and resulted from evolutionary pressure (Figure 6.c). The basic idea behind this method is that any randomization of orthologous sequences of one protein will disrupt the phylogeny and destroy and evolutionary pressure between the two proteins (see Figure 6.b). The statistical significance of the heterodimerization is therefore computed as the total number of randomized ECs with more significant scores than the original, 1 EC, divided by the total number of randomized phylogenies, 100, leading to an assessed p-value of 0.01.

**Figure 6:**
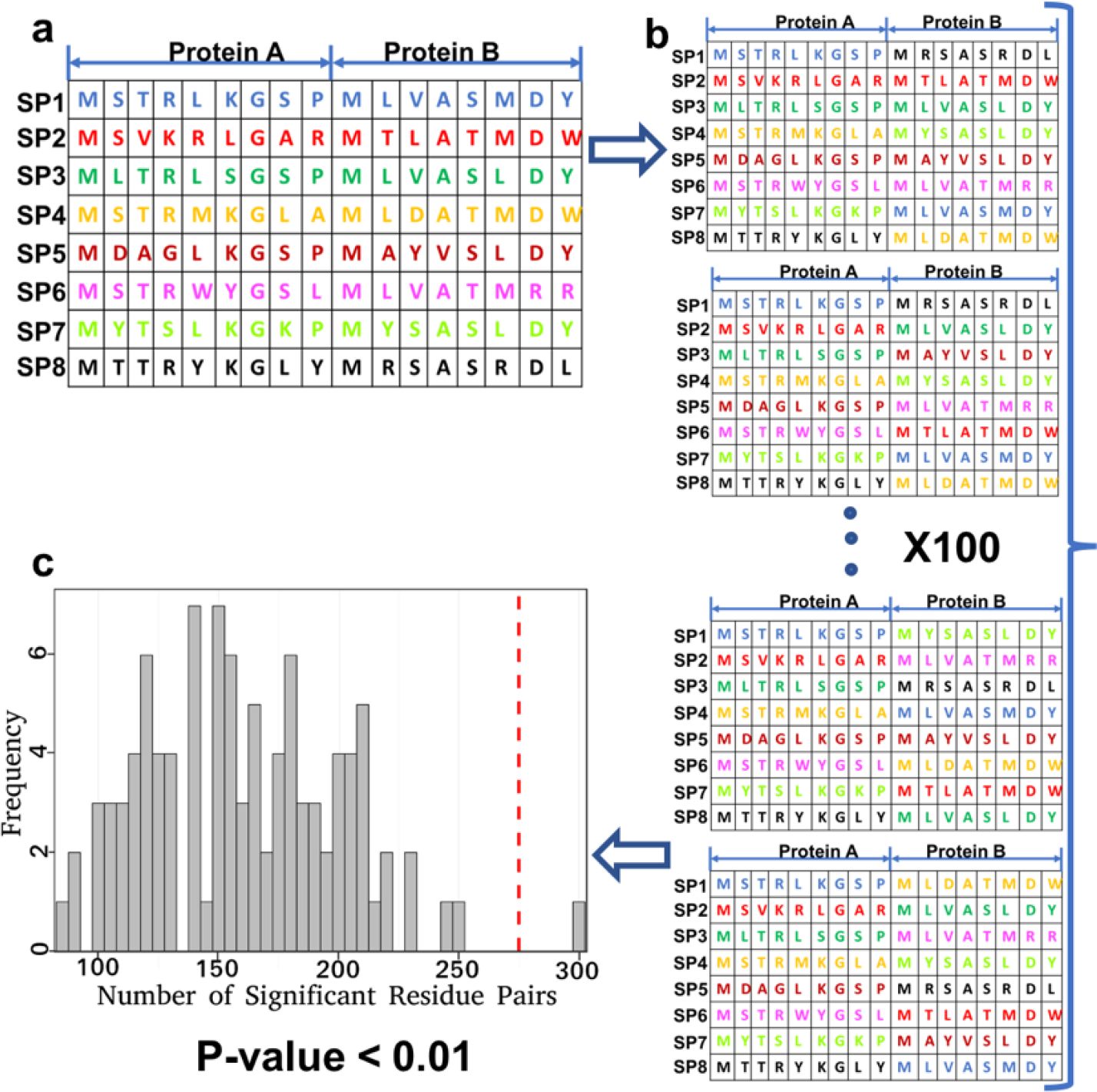
Randomized phylogeny to assess the statistical significance of the heterodimerization. We randomized MSA of the 5-HT_2C_R while keeping the 5-HT_2A_R. Estimate the No of residue signal > the minimal of 5% of the original signal and found that only one randomized ECs has more significant ECs leading to a p-value < 0.01 (see text for description)

**Figure 7:**
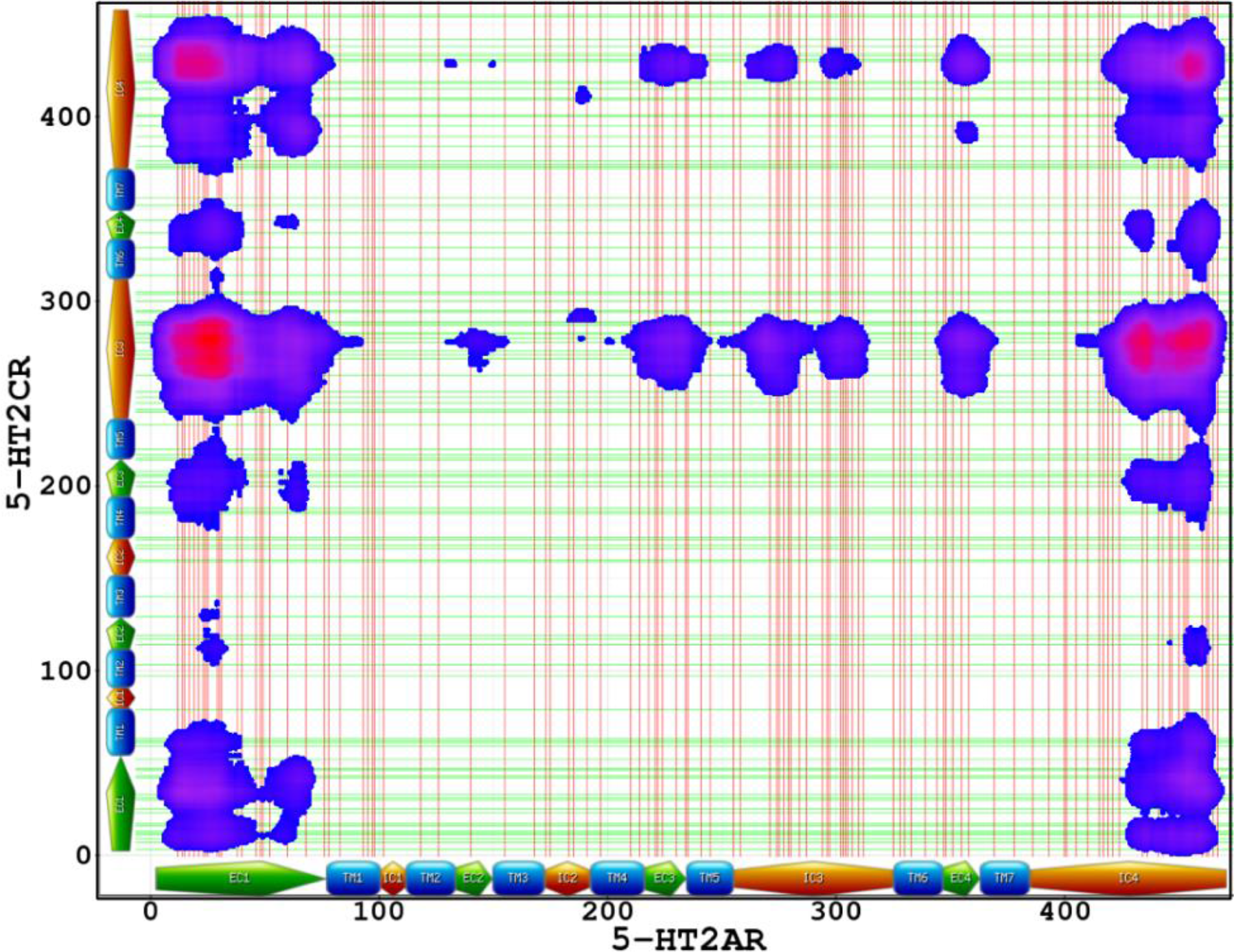
SNPs positions at the interacting peptides. SNPs positions in 5-HT_2A_R (red lines) and 5-HT_2C_R (green lines) are plotted and compared to most likely interacting peptides. The density of SNPs appears to correlate with predicted interacting peptides.

### Polymorphisms associated with inferred contacts

It has been hypothesized that the 5-HT_2A_R/5-HT_2C_R interaction plays a role in substance use disorders. There is significant variation in individual propensity to addiction disorders, attributed to genetic variants, including polymorphisms in receptor proteins (Comings, Comings et al. 1991, Bond, LaForge et al. 1998, Szeto, Tang et al. 2001, Wang, Quillan et al. 2001, Oslin, Berrettini et al. 2003, Kreek, Bart et al. 2005, Liu, Drgon et al. 2006, van den Wildenberg, Wiers et al. 2007, Levran, Peles et al. 2014, Levran, Peles et al. 2015). Also, systematic difference exists between male and female subjects (Brady and Randall 1999, Perkins and Scott 2008). It is possible that some of these differences might be explained by genetic variants affecting the interaction between 5-HT_2A_R and 5-HT_2C_R, specifically by polymorphisms in receptor coding sequences that are translated to residues involved in the interaction. Also, the statistical differences observed between the sexes may be caused by different propagation of genotypic variants to phenotype in males vs females in the case of the X-linked 5-HT_2C_R gene.

To identify genetic polymorphisms of 5-HT_2A_R and 5-HT_2C_R that may affect the interaction, we queried the *NCBI* database of genetic variation (Sherry, Ward et al. 2001) and extracted common variants coinciding with the coding sequences of HTR2A and of HTR2C. Next, we selected the SNPs that are located on the interacting residues. We considered the affected residues when convolved ECs was greater than 0.01 (corresponding to the top 0.1% of the map). The list of the most studied SNPs is in Table 3 and the full list is provided as supplementary material (Supplementary data File SDF3).

**Table 1:** Receptor properties. The significant length of the complex (total length of the cMSA / # of sequences) was computed to be 0.25, lower than the 0.7 previously suggested for DCA analysis in single proteins. As explained in the text, the convolution of local ECs scores and secondary structure helps highlight the interacting peptides even with low significant length.

|  | Protein length | # individual sequences | cMSA | Sig. length | Sequence coverage |
| --- | --- | --- | --- | --- | --- |
| 5-HT<sub>2C</sub>R | 458 AA | 244 | 231 | 0.25 | 55 – 90% |
| 5-HT<sub>2A</sub>R | 471 AA | 262 |  |  |  |

**Table 2:**
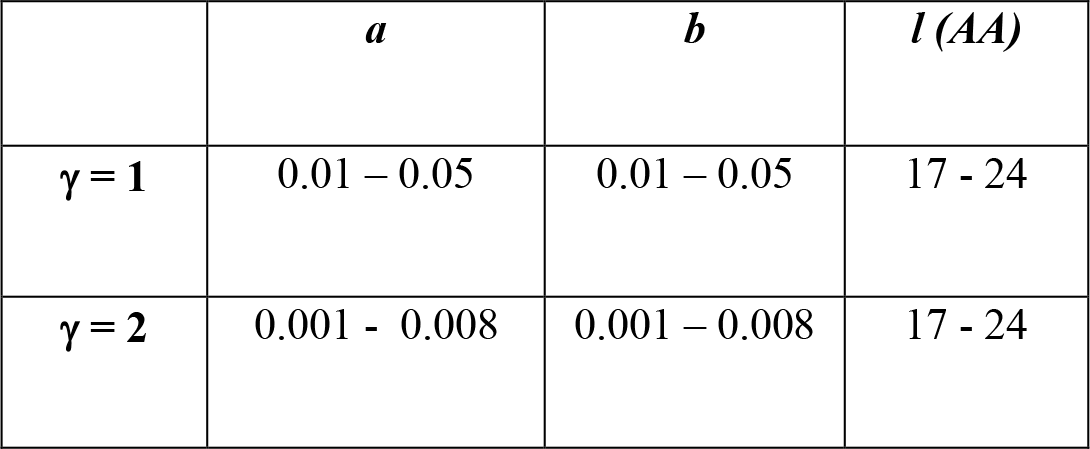
Optimized values of the Gaussian Convolution: Values were optimized using serotonin proteins and GPCRs. a and b are the optimized variances of the Gaussian Kernel, l is the average length of the interacting peptides and γ = 1 if the residues belong to the same secondary structure, if not γ = 2.

|  | *a* | *b* | *l* (AA) |
| --- | --- | --- | --- |
| $\gamma = 1$ | 0.01 – 0.05 | 0.01 – 0.05 | 17 - 24 |
| $\gamma = 2$ | 0.001 - 0.008 | 0.001 – 0.008 | 17 - 24 |

**Table 3:** Significant SNPs at interacting peptides of 5-HT_2A_R – 5-HT_2C_R: full list of SNPs located at the interacting peptides is provided in supplementary data file SDF3.

| *SNPs ID* | *SNP in* | *5-HT2AR position* | *5-HTR2A AA* | *5-HTR2A Domain* | *5-HT2CR position* | *5-HTR2C AA* | *5-HT2CR Domain* | *Convolved ECs* |
| --- | --- | --- | --- | --- | --- | --- | --- | --- |
| rs6314 | 5-HT2AR | 452 | H | IC4 | 278 | A | IC3 | 0.049 |
| rs769223465 | 5-HT2AR | 467 | K | IC4 | 286 | A | IC3 | 0.030 |
| rs773466572 | 5-HT2AR | 78 | A | TM1 | 278 | A | IC3 | 0.025 |
| rs749275325 | 5-HT2AR | 463 | G | IC4 | 280 | P | IC3 | 0.043 |
| rs781643370 | 5-HT2AR | 214 | G | TM4 | 279 | N | IC3 | 0.023 |
| rs75634836 | 5-HT2AR | 241 | V | TM5 | 428 | S | IC4 | 0.022 |
| rs6318 | 5-HT2CR | 25 | T | EC1 | 23 | C | EC1 | 0.025 |
| rs143566182 | 5-HT2CR | 453 | S | IC4 | 42 | D | EC1 | 0.034 |
| rs781896215 | 5-HT2CR | 457 | S | IC4 | 59 | S | TM1 | 0.027 |
| rs144618891 | 5-HT2CR | 454 | E | IC4 | 188 | P | TM4 | 0.022 |
| rs144618891 | 5-HT2CR | 457 | S | IC4 | 188 | P | TM4 | 0.022 |
| rs368641796 | 5-HT2CR | 457 | S | IC4 | 214 | F | TM5 | 0.022 |
| rs782627259 | 5-HT2CR | 460 | N | IC4 | 323 | M | TM6 | 0.023 |

Several SNPs found to be involved in the dimerization of 5-HT_2A_R and 5-HT_2C_R have been implicated in neuropsychiatric diseases and aggressive behavior (impulsivity, hostility). On the HTR2C gene, rs6318 also known as Cys23Ser or 68G>C, is implicated in eating and anxiety disorders, obesity, depression, bipolar disorder, schizophrenia, and cocaine use disorder. On the HTR2A gene, rs6314, also known as His452Ty or C1354>T, is associated with depression and Cocaine use disorder.

An interesting observation is that polymorphisms in the genes coding for the HTR2A and HTR2C proteins are more prevalent in the intra- and extracellular domains (75 SNPs in 5-HT_2C_R, on average 26 SNPs/100AA; 93 SNPs in 5-HT_2A_R, on average 30 SNPs/100aa) than in the transmembrane domains (31 SNPs in 5-HT_2C_R, on average 19 SNPs/100AA; 25 SNPs in 5-HT_2A_R, on average 16 SNPS/100aa). Therefore, polymorphisms of clinical relevance affecting the function (including dimerization) of these proteins and potentially related to propensity toward substance use disorders, are more likely to exist in the IC and EC domains, even if interactions also exist in the TM regions of the two proteins. Intriguingly, this pattern is similar to the coincidence between interspecific and intraspecific variation previously observed in mRNA processing regions (Castle 2011), and in disease-associated genes (Miller and Kumar 2001).

### Coexpression of HTR_2A_ and HTR_2C_ in baboon circadian cycle

Analysis of gene expression data is a valuable tool for functional analysis of genes and proteins. Specifically, products of co-expressed genes are often involved in direct protein-protein interaction (Lee, Hsu et al. 2004). Recently, a large timecourse study of baboon circadian circle has been published that covers circadian transcriptome in 64 different tissues, including 22 different brain regions, sampled every 2 h over 24 h.

We computed the coexpression between the circadian profiles of the 5-HT_2A_R and 5-HT_2C_R receptors as the Pearson Correlation coefficient between the two transcripts (Kudlicki, Rowicka et al. 2007). We find that for most brain regions the correlation is positive. To assess the significance of the positive correlation, we calculated the fraction of named genes that have correlation with 5-HT_2A_R lower than 5-HT_2C_R and the fraction of named genes that have correlation with 5-HT_2C_R lower than 5-HT_2A_R. The results are summarized in Table 4. While this result does not prove that 5-HT_2A_R and 5-HT_2C_R are involved in a direct interaction, it indicates in which tissues such functional contact may be taking place. One notable finding is that the correlation is especially high and statistically significant in the prefrontal cortex, a brain region associated with forming addiction (Goldstein and Volkow 2011). The result is consistent with the hypothesis that the 5-HT_2A_R/5-HT_2C_R interaction indeed takes place and that it may play a role in substance abuse disorders (Anastasio, Gilbertson et al. 2013, Anastasio, Liu et al. 2014, Anastasio, Stutz et al. 2014, McAllister 2016).

**Table 4:** Gene expressions of HTR_2A_ and HTR_2C_ are correlated in baboon circadian cycle. PCC: Pearson Correlation Coefficient; **FRPG – Random**: Fraction of Random Pairs of named Genes with lower PCC in the same dataset (based on 10000 random pairs); **FNG - HTR_2A_**: Fraction of Named Genes that have lower PCC with HTR_2A_ in the same dataset; **FNG - HTR_2C_**: fraction of named genes that have lower PCC with HTR_2C_ in the same dataset.

| *TISSUE* | *PCC* | *FRPG - Random* | *FNG - HTR2A* | *FNG - HTR2C* | *TISSUE NAME* |
| --- | --- | --- | --- | --- | --- |
| VIC | -0.4740 | 0.0871 | 0.0390 | 0.2852 | Visual cortex |
| PUT | -0.2670 | 0.1822 | 0.1420 | 0.2591 | Putamen |
| HAB | -0.1570 | 0.2773 | 0.3048 | 0.3273 | Habenula |
| SCN | -0.1040 | 0.2993 | 0.2258 | 0.4452 | Suprachiasmatic nuclei |
| CER | -0.0020 | 0.3133 | 0.5560 | 0.3693 | Cerebellum |
| LH | 0.0020 | 0.6056 | 0.5847 | 0.6835 | Lateral hypothalamus |
| PON | 0.0270 | 0.6426 | 0.5594 | 0.7179 | Pons |
| VMH | 0.0460 | 0.6386 | 0.5765 | 0.5239 | Ventromedial hypothalamus |
| ARC | 0.0590 | 0.6076 | 0.5665 | 0.5481 | Arcuate nucleus |
| DMH | 0.0940 | 0.6757 | 0.5685 | 0.6707 | Dorsomedial hypothalamus |
| MGP | 0.2150 | 0.7568 | 0.7730 | 0.8283 | Medial globus pallidus |
| LGP | 0.2470 | 0.7928 | 0.7237 | 0.8563 | Lateral globus pallidus |

**Table 4: Gene expressions of HTR2A and HTR2C are correlated in baboon circadian cycle.**
|  |  |  |  |  |  |
| --- | --- | --- | --- | --- | --- |
| PVN | 0.2880 | 0.7648 | 0.8034 | 0.6770 | Paraventricular nuclei |
| SUN | 0.3040 | 0.7998 | 0.9069 | 0.8041 | Substantia nigra |
| PRA | 0.3050 | 0.8038 | 0.8143 | 0.7510 | Preoptic area |
| HIP | 0.3570 | 0.8228 | 0.7940 | 0.8293 | Hippocampus |
| AMY | 0.4920 | 0.7778 | 0.9025 | 0.9463 | Amygdala |
| ONH | 0.5860 | 0.8939 | 0.8970 | 0.9507 | Optic nerve head |
| SON s | 0.6250 | 0.9489 | 0.9138 | 0.9200 | Supraoptic nucleus |
| OLB | 0.6660 | 0.9449 | 0.8654 | 0.9791 | Olfactory bulb |
| PRC | 0.6880 | 0.9509 | 0.9851 | 0.9747 | Prefrontal cortex |
| MMB | 0.7490 | 0.9640 | 0.9771 | 0.9590 | Mammillary body |
| THA | 0.7780 | 0.9770 | 0.9742 | 0.9363 | Thalamus |

## MATERIALS AND METHODS

### Multiple Sequence Alignment and Evolutionary Scores

Evolutionary coupling analysis depends on a large collection of genes orthologous to each of the genes whose interaction is under investigation; the orthologs of both genes have to come from the same set of species. We obtained 241 and 235 orthologous sequences of 5-HT_2A_R and 5-HT_2C_R respectively by querying *GENBANK* (Benson, Lipman et al. 1993) and running genome-wide *tblastn* (Altschul, Gish et al. 1990) on genomes not present in *GENBANK*. The orthologous sequences were joined by species and aligned using *Clustal-Ω* (Sievers, Wilm et al. 2011) with manually introduced corrections (based on domain annotation) to generate the concatenated Multiple Sequence Alignments (cMSA) of 5-HT_2A_R/5-HT_2C_R in 231 vertebrates. The list of species is provided as supplementary data file (SDF1). The alignment is provided as supplementary Figure S1.

We analyzed evolutionary coupling using Direct Coupling Analysis (DCA) algorithm (Morcos, Pagnani et al. 2011). DCA was chosen among Evolutionary Coupling (EC) methods because of its better ability to model the whole data set rather than individual pairs of residues, helping to distinguish between direct functional residue interactions and correlations resulting from indirect interactions that are given a lower weight in DCA. The primary advantage of DCA lies in this ability to assign higher scores to direct correlations than to indirect ones, which is achieved through computing direct statistical couplings for each pair of positions and an appropriate sampling correction. The pseudo-likelihood maximization Direct-Coupling Analysis (*plmDCA*) variant of the algorithm (Ekeberg, Hartonen et al. 2014) that we utilize has a lower computational cost than alternative approaches. Options for *plmDCA* were set as follows: Optimization method (conjugate gradient, cg); Sequence of interest (human, use first member as sequence of interest); and other parameters as default.

### DCA post-processing: convolution of Evolutionary Coupling scores (ECs): 2D Gaussian filter applied to DCA maps

DCA-based methods have been shown to be reliable in predicting native contacts and have long been used to reconstruct the 3D structure of proteins. However, the ECs scores, as provided by these methods, are noisy and contain a large number of False Positives (FP), especially for large proteins, such as 5-HT_2A_R and 5-HT_2C_R. Such FPs, due to random correlations between evolutionary changes, appear as isolated peaks in the DCA maps. To account for the contribution of neighboring residues in the formation of interfaces, we propose here to use a Gaussian convolution with a kernel constructed on the structural information of both proteins. Convolution or deconvolution methods have been successfully used in the past to extract the signal in various processes (Kudlicki, Plewa et al. 1996, Rowicka, Kudlicki et al. 2007, Fongang and Kudlicki 2013, Homouz, Chen et al. 2015, Fongang and Kudlicki 2016) in which the background noise or experimental artifacts overshadow the true signal.

Our global approach is outlined in Figure 1: For two studied proteins, a cMSA is constructed from orthologous sequences and ECs are computed using the *plmDCA* algorithm as previously described. For each pair of residues *i,j*, a 2D Gaussian convolution, centered on (*i,j*) with different variances is applied resulting in convolved ECs that depend on the local structure of peptides. ECs provided by the DCA algorithm is a M by N matrix where M and N are the lengths of the two interacting proteins. Each point of the matrix, *P*_*i,j*_, represents the Evolutionary Score of residues *i* and *j* of the two proteins. The convolved DCA signal is defined as:

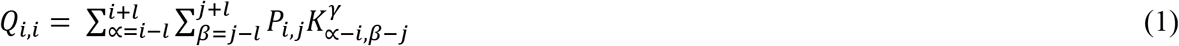

where 2*l* + *1 is* the number of residues in the neighborhood and 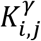 is a kernel function adjusted to the secondary structure. The biological interpretation of *l* is that it represents the average length of the stretch of protein sequence involved in the interaction. In the present study, the kernel 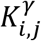 is a Gaussian function

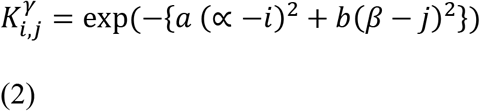

with *a* and *b* the normalized variances and γ denotes the dependence to the secondary structure of the protein (γ=1 if the same type of structure in both proteins, γ=2 otherwise; optimized values of a b for both situations are given in Table 2). The general idea behind the local convolution of ECs signal is depicted in Figure 1: Residues with the same secondary structure (alpha helix, beta barrel or coil) but less significant overall ECs have greater chance to be in contact that isolated residues with stronger ECs. The final convolved ECs depends on the variances *a* and *b* as well as *l*, the total number of amino-acids (AA) at the interface. Optimized values of the parameters *a* and *b*, estimated on the present serotonin receptors and several other GPCRs are listed in Table 2. We have estimated the parameter *l* = *21 AA* for the 5-HT_2A_R-5HT_2C_R heterodimer (see Results and Discussion section). The Gaussian convolution depends on the accurate prediction of the structural information of individual protein. Here, we predicted the secondary structure of 5- HT_2A_R and 5-HT_2C_R using stand-alone versions of PSIPRED (McGuffin, Bryson et al. 2000), with manual correction for consistency. This bioinformatics tool has been shown to accurately predict protein structure information based on sequences and we have verified that predicted secondary structures, in the case of 5-HT_2A_R and 5-HT_2C_R agree with known properties (GPCRs domain architecture). The secondary structures were checked for consistency and corrected where required using our own program written in R.

### Single Nucleotide Polymorphisms (SNPs)

Human SNPs localized in the coding sequence of 5-HT_2A_R and 5-HT_2C_R were retrieved from the NCBI SNPs database (dbSNP) (Sherry, Ward et al. 2001) and analyzed in terms of coevolution and phenotypic annotation (see Results).

### Coexpression during baboon circadian rhythm

To assess coexpression of the receptor genes in primate circadian cycle, we have downloaded transcriptome data from the diurnal transcriptome atlas of Baboon (Papio Anubis) across all major neural and peripheral tissues (64 tissues over 12 time points), available at GEO under accession number GSE98965 (Mure, Le et al. 2018). Correlation between the genes coding for 5-HT_2A_R and 5-HT_2C_R has been computed over time. To assess the significance of co-expression the correlations were compared with correlations between 10,000 randomly chosen pairs of named genes, as well as between the serotonin receptors and all other named genes in the dataset.

## CONCLUSIONS

Whether 5-HT_2A_R and 5-HT_2C_R form a functional heterodimer has been a subject of debate over the last decade, while recent evidence strongly points towards the heterodimer forming at least in some conditions. The two receptors are particularly important as they are involved in substances abuse and may be used to design drugs targeting substance addiction. Several experimental evidences point towards the formation of a 5HT_2A_R/5-HT_2C_R dimer in prefrontal cortex of rats and other model animals, but where and how this heterodimerization is formed remains elusive. Using computational methods and the co-evolution principle of functional residues of proteins, we concluded that the 5-HT_2A_R and 5-HT_2C_R form a functional heterodimer with interacting peptides either on the N-terminal domain, 3rd intracellular loop or the C-terminal domain of the receptors. Although significant interactions are highlighted using the local deconvolution method, signal also remains that points to several interactions involving physically distant domains. A possible explanation is the existence of allosteric interactions linking residues within a protein across its domains. Such allosteric interactions are crucial to normal function of transmembrane receptor proteins.

Traditional DCA methods work of large number of sequences (N/L > 0.7), but here we show that if one is only interested in predicting the most likely peptides, a local convolution of ECs signals with the kernel adjusted to the secondary structure element is an efficient post-processing method. Moreover, the post-processing opens the door to the investigation of the heterodimerization in larger multi-domain proteins. We assessed the statistical significance of the 5-HT_2A_R/5-HT_2C_R dimerization using the randomized phylogeny and concluded that the p-value (probability that the observed interaction is due to noise) < 0.01. As it has been reported that the Single Nucleotide Polymorphisms (SNPs) play a significant role in differential propensity to substance abuse disorders, we determined the SNPs and found that a significant number of SNPs are located at the binding interfaces. Finally, an analysis of coexpression in the recently published primate transcription atlas suggests the prefrontal cortex as one possible tissue where the interaction between 5-HT_2A_R/5-HT_2C_R may be functional.

## Supporting information

Supplementary data file SDF3

Supplementary Data File SDF1

Supplementary Data File SDF2

## ACKNOWLEDGMENTS

This study was conducted with the support by NIH grants P50 DA033935. R01 GM112131, the Institute for Translational Sciences, supported in part by a Clinical and Translational Science Award (UL1TR000071) from the National Center for Advancing Translational Sciences, and the Center for Addiction Research at the University of Texas Medical Branch. The authors are very grateful to Noelle C. Anastasio, Scott R. Gilbertson, F. Gerard Moeller for insightful discussions.

## Authors contributions

AK and BF designed research; BF and AK wrote computer code; BF prepared illustrations; BF calculated coevolution maps and performed statistical analysis; AK analyzed expression profiles across tissues; BF, MR and AK interpreted results; KC integrated results with experimental data; BF, MR, AK and KC wrote the paper.

## SUPORTING INFORMATION

Supplementary data File SDF1: cMSA of 5-HT_2A_R – 5-HT_2C_R: list of species used to construct the concatenated multiple sequence alignments. cMSA is built by joining orthologs 5-HT_2A_R and 5-HT_2C_R from the same species.

Supplementary Figure SF1: cMSA of 5-HT_2A_R – 5-HT_2C_R: cMSA is built by joining orthologs 5-HT_2A_R and 5-HT_2C_R from the same species. Transmembrane domains are highly conserved (% identity > 90) between species.

Supplementary Data File SDF2: Convolved ECs of 5-HT_2A_R – 5-HT_2C_R:

Supplementary data file SDF3: list of all SNPs:

