## Supplementary figures and images for "Protein Co-Evolution Strategies Detect Predicted Functional Interaction Between the Serotonin 5-HT_2A_ and 5-HT_2C_ Receptors"

### Supplementary Figure SF1

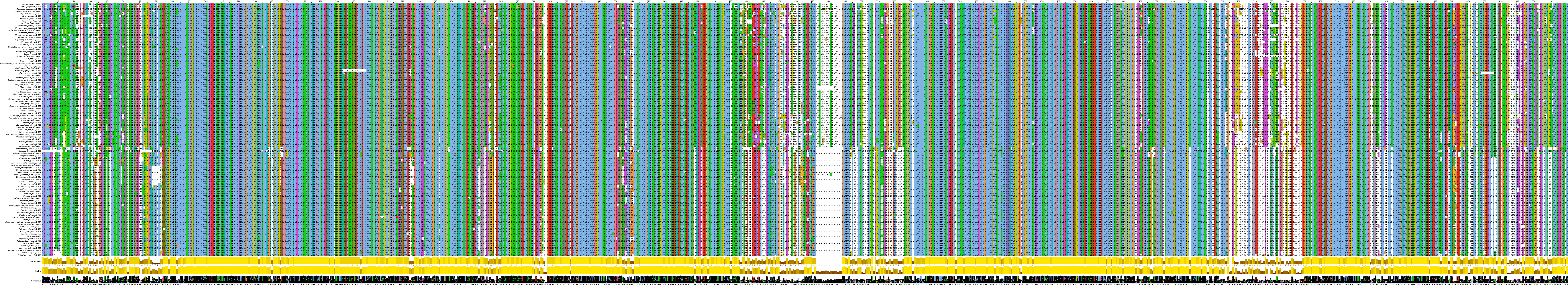
